# Apical3DTip: Elliptic Cross-section-based Reconstruction for the Embryo Initial Cell of Arabidopsis

**DOI:** 10.64898/2026.06.25.734685

**Authors:** Tomonobu Nonoyama, Zichen Kang, Yuga Hanaki, Yoshinobu Itagaki, Hikari Matsumoto, Yusuke Kimata, Satoru Tsugawa, Minako Ueda

## Abstract

**Background:** Cell geometry plays a central role in determining division orientation and body axis formation during early embryogenesis in *Arabidopsis thaliana*. However, quantitative analysis of dynamic three-dimensional (3D) morphology remains challenging because live-imaging studies often rely on two-dimensional (2D) projections, while existing 3D reconstruction approaches, including mesh-based methods, often lose the original orientation information relative to the ovule and require labor-intensive mesh correction. In addition, embryo positional fluctuation caused by floating in liquid medium and continuous growth makes it difficult to analyze temporal morphological changes within a common coordinate system.

**Results:** We developed a robust framework for quantitative 3D and four-dimensional (4D; 3D + time) analysis of embryo initial cell (apical cell) morphology. The method first establishes a standardized 3D coordinate system by normalizing cell orientation based on the bottom plane and the optical axis of the observation. Cell morphology is then reconstructed through ellipse-based approximation of serial cross-sections extracted from stacked imaging data, enabling accurate geometric characterization without the need for complex surface mesh reconstruction. To evaluate shape anisotropy, we quantified the apical cell shape in 3D. The framework further supports the characterization of volumetric features of subsequent division, providing a basis for quantifying 3D embryogenesis.

**Conclusion:** Our framework provides a simple and noise-reduced approach for quantitative analysis of living cell morphology in 3D. We named the integrated method of combining coordinate normalization with elliptical cross-section-based reconstruction Apical3DTip. This method enables consistent comparison of cell shapes without extensive manual corrections. The method overcomes key limitations of 2D projection-based and mesh-dependent analyses and offers a practical platform for quantifying cell shape and daughter cell shapes in 3D. More broadly, it provides a quantitative foundation for exploring the relationship between cell geometry, morphodynamics, and developmental patterning in living plant embryos.

## Background

Three-dimensional (3D) reconstructions of fixed embryos of *Arabidopsis thaliana* have shown that cell division patterns follow predictable geometric rules that are tightly coupled to body axis formation [1, 2]. By segmenting cells in 3D and modelling division planes, these studies demonstrated that divisions tend to minimize surface area or follow shortest-wall principles proposed in the early studies [3, 4, 5], linking cell shape directly to division orientation. In the early embryo, such geometrically constrained divisions establish tissue patterning and axis formation [6]. Importantly, computational simulations confirmed that even slight deviations in cell geometry could alter division orientation, emphasizing that geometry is not merely descriptive but instructive [1]. These findings support a framework in which intrinsic physical constraints guide developmental patterning, suggesting that morphogenesis emerges from the integration of cell shape, possibly mechanical forces, and division rules. Thus, 3D analysis has been essential in demonstrating that axis formation is not solely genetically preprogrammed but is also a consequence of geometric and biophysical principles acting at the cellular level.

Recent studies based on live-cell imaging showed that even slight morphological changes can also significantly influence the division orientation in the apical cells [7]. This highlights the importance of analyzing cells as 3D structures, however, current live-imaging studies of embryogenesis remain limited to 2D analyses, mainly due to the following three constraints. First, during live-cell imaging, embryos are cultured in liquid medium and are not rigidly fixed, causing substantial image fluctuations that require continuous drift correction [6]. Second, laser intensity must be kept low to avoid photodamage during long-term observation, often resulting in weak signals from deeper regions and incomplete visualization of cell boundaries. Finally, these limitations make it difficult not only to reconstruct accurate 3D cell geometries but also to maintain a common coordinate system that preserves embryo orientation within the ovule, imaging direction, and growth-related positional changes over time [6]. As a result, quantification of the dynamics of morphology in apical cells and subsequent embryos in 3D remains unresolved. While a promising tool MorphoGraphX for 3D cell segmentation provides a precise 3D morphology [8], it sometimes requires manual corrections of mesh, especially when the contours are disconnected due to weak signals, and thus is not very suitable for tracking numerous time lapse image sequences. Also, cell fluctuation in liquid media needs to be analyzed in a well-defined coordinate system. Therefore, it requires a precise, simple, and robust method of 4D morphodynamics (3D + time) in this field.

In this study, we developed a 3D coordinate normalization framework (Apical3DTip) based on the apical cell bottom, which is a relatively flat boundary with the adjacent basal cell, to establish a common spatial reference for apical cells. This framework corrects positional fluctuations caused by sample movement while preserving biologically relevant information, including embryo orientation within the ovule and imaging geometry. After normalization, we applied elliptical cross-section-based reconstruction to cell imaging data, enabling interpolation of incompletely detected cell boundaries in deep regions and reducing the need for labor-intensive mesh correction. Directional morphology was then quantified by bounding-box analysis in all circumferential orientations. Further analyses of divided apical cells demonstrated that our method robustly captured 3D morphological traits around the cell axis of subsequent embryogenesis.

## Results

### 1. 3D coordinate normalization based on bottom plane of the apical cell

To eliminate positional deviations caused by sample drift and growth and to establish a standardized coordinate system, we used a brighter point cloud (bottom plane), which is formed by the zygotic division. Based on this bottom plane, we developed a 3D normalization method as follows (see details in Methods). First, *Z*-stack two-photon excitation microscopy (2PEM) images showing plasma membrane (PM) as the apical cell outline were obtained from the raw (*X, Y, Z*) coordinate system, where *Z*-axis corresponds to optical axis (Fig. 1a). Then, a 3D spatial coordinate is constructed by assigning each voxel a position in space along with intensity value (Fig. 1b). This step transforms the stacked image data into a structured volumetric representation for quantitative analysis. Second, an appropriate intensity threshold is applied to isolate the bottom plane by discarding surrounding components (Fig. 1c). Third, the spatially averaged normal vector of the division plane is calculated (Fig. 1d). This provides a robust representation of its overall geometry. Fourth, the original bottom plane undergoes *Z*-axis rotation which transforms the coordinate system into (*X*_1_, *Y*_1_, *Z*_1_) (Fig. 1e), and *X*_1_ -axis rotation which transforms the coordinate system into *X*_2_ = *x, Y*_2_ = *y, Z*_2_ = *z*) (Fig. 1f).

**Fig. 1.**
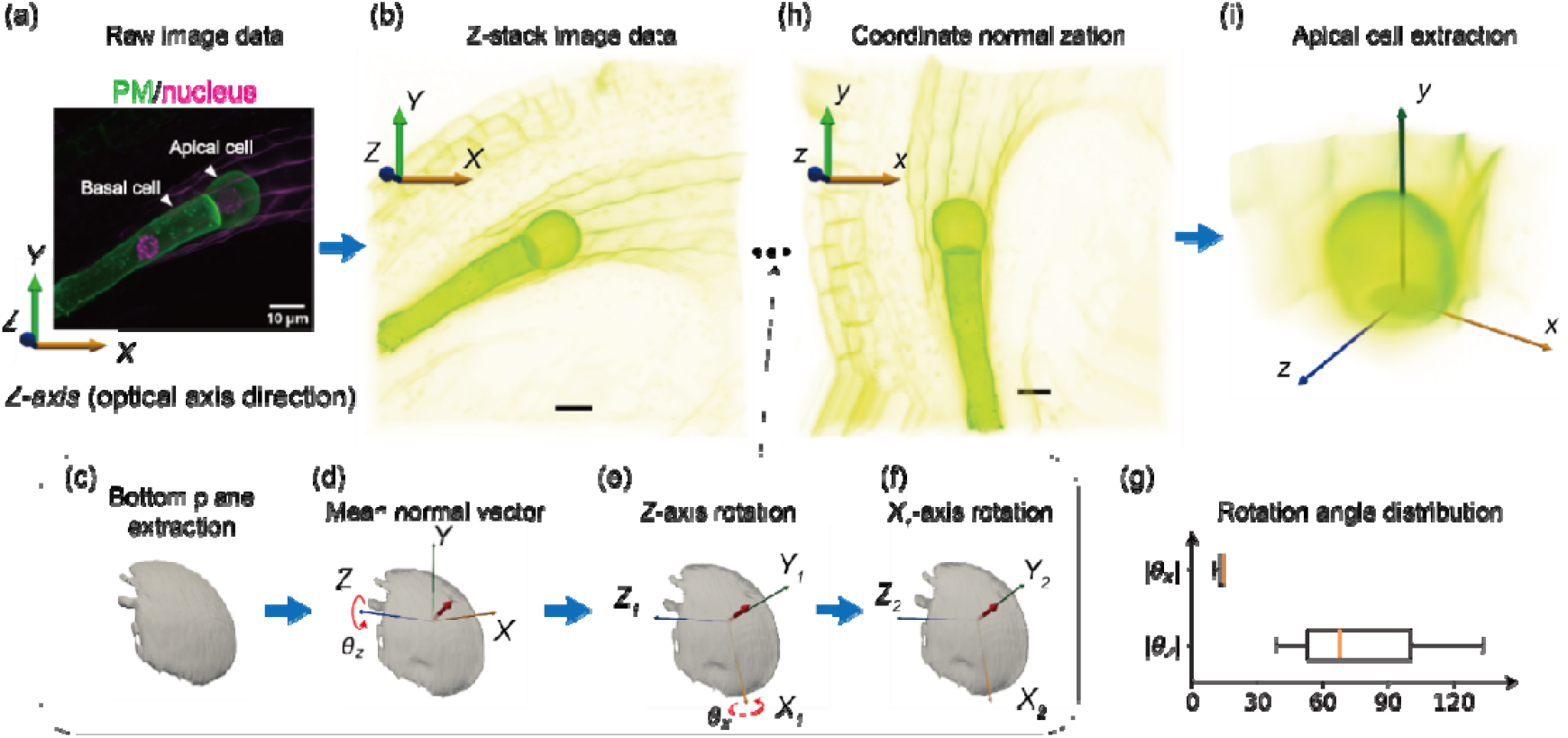
Workflow of 3D coordinate normalization for apical cell analysis. (a) The raw ***Z***-stack 2PEM image data of the divided zygote expressing the PM and nucleus marker. Two daughter cells, an apical cell and a basal cell, are detected. (b) ***Z***-stack image data with the original coordinate system (***X, Y, Z***). (c) Identification of the bottom plane using a high intensity threshold (>99%). (d) Refinement of the extracted plane by eliminating isolated noise voxels and selecting the largest connected component. The mean normal vector is denoted by the red vector. (e) ***Z***-axis rotation of the plane keeping ***Z***-stack information after the rotation. (f) ***X***-axis rotation with a rotation angle. (g) The range of ***Z***-axis and ***X***-axis rotation angle ***θ***_***Z***_ and ***θ***_***X***_. (h) Alignment of the entire cell into a new coordinate system (***x, y, z***) by the two rotations so that the mean normal vector is oriented along the positive ***y***-axis and the bottom plane becomes parallel to the ***xy***-plane (3D coordinate normalization). (i) Apical cell extraction in the new coordinate system.

In most cases, the elongated embryo is located within the relatively flat seed and lies approximately perpendicular to the gravity direction during imaging, as illustrated in Fig. 1a. As a result, the *X*_1_-axis rotation angle (*θ*_*X*_) required for coordinate normalization should remain small, and the corrected z-axis should be nearly aligned with the original optical axis (*Z*-axis) of the imaging system. Consistent with this expectation, *θ*_*X*_ was less than 15 degrees for all three samples, suggesting only a slight tilt of the original imaging data around the *X*-axis (Fig. 1g). In contrast, the in-plane orientation of the seed is essentially random within the plane. Accordingly, the *Z*-axis rotation angle (*θ*_*Z*_), which does not affect the *Z*-stack intensity, exhibited relatively large values ranging from 60 to 100 degrees across the three samples (Fig. 1g). Finally, the voxel data of the entire cell is aligned through these rotations making the normal direction *y*-axis, which is termed 3D coordinate normalization (Fig. 1h). The transformation is performed so that the computed average normal vector points in the positive *y*-direction (i.e., the apical-basal axis of the embryo), and the bottom plane becomes parallel to the *xz*-plane (i.e., the radial axis), standardizing the orientation of the apical cell for consistent comparison and analysis (Fig. 1i).

### 2. Quantitative reconstruction of 3D cell morphology via elliptical cross-section analysis

Next, we developed a method to accurately reconstruct cell contours even from the weak signals in deep regions. After 3D coordinate normalization, the cell-aligned along the new *y*-axis, is analyzed by extracting cross-sectional contours on planes parallel to the *xy*-plane (Fig. 2a). For each *z*-slice, the contour is detected using radial scanning centered at the intensity centroid (Fig. 2b). Along each radial line, the outermost pixel exceeding a predefined threshold is selected as a sample point. When multiple clusters of such points appear along a scan line, the cluster with the largest number of points is chosen as the representative set, while smaller clusters are discarded to avoid noise and artifacts (Fig. 2c). The resulting representative point set is then approximated by an ellipse, providing a simplified geometric description of the cell cross-section (Fig. 2d). The accuracy of this approximation is quantitatively evaluated using the root mean square error (*RMSE*) relative to the major axis (*r*_*a*_), ensuring that the fitted ellipse adequately reflects the original intensity-based contour (Fig. 2e). Finally, the major and minor axes of the ellipse, denoted as *r*_*a*_ and *r*_*b*_, are smoothed along the *y*-axis using Lowess regression in python (see Methods) (Fig. 2f). This smoothing step reduces slice-to-slice variability introduced by independent ellipse fitting, thereby correcting fluctuations in the reconstructed 3D shape and yielding a more consistent representation of the cell morphology (Fig. 2f). We called all the processes elliptic cross-section-based reconstruction and wrote some details in Methods. By integrating this method with the 3D coordinate normalization described above, we were able to construct 3D shapes of living apical cells, and we named this integrated method Apical3DTip.

**Fig. 2.**
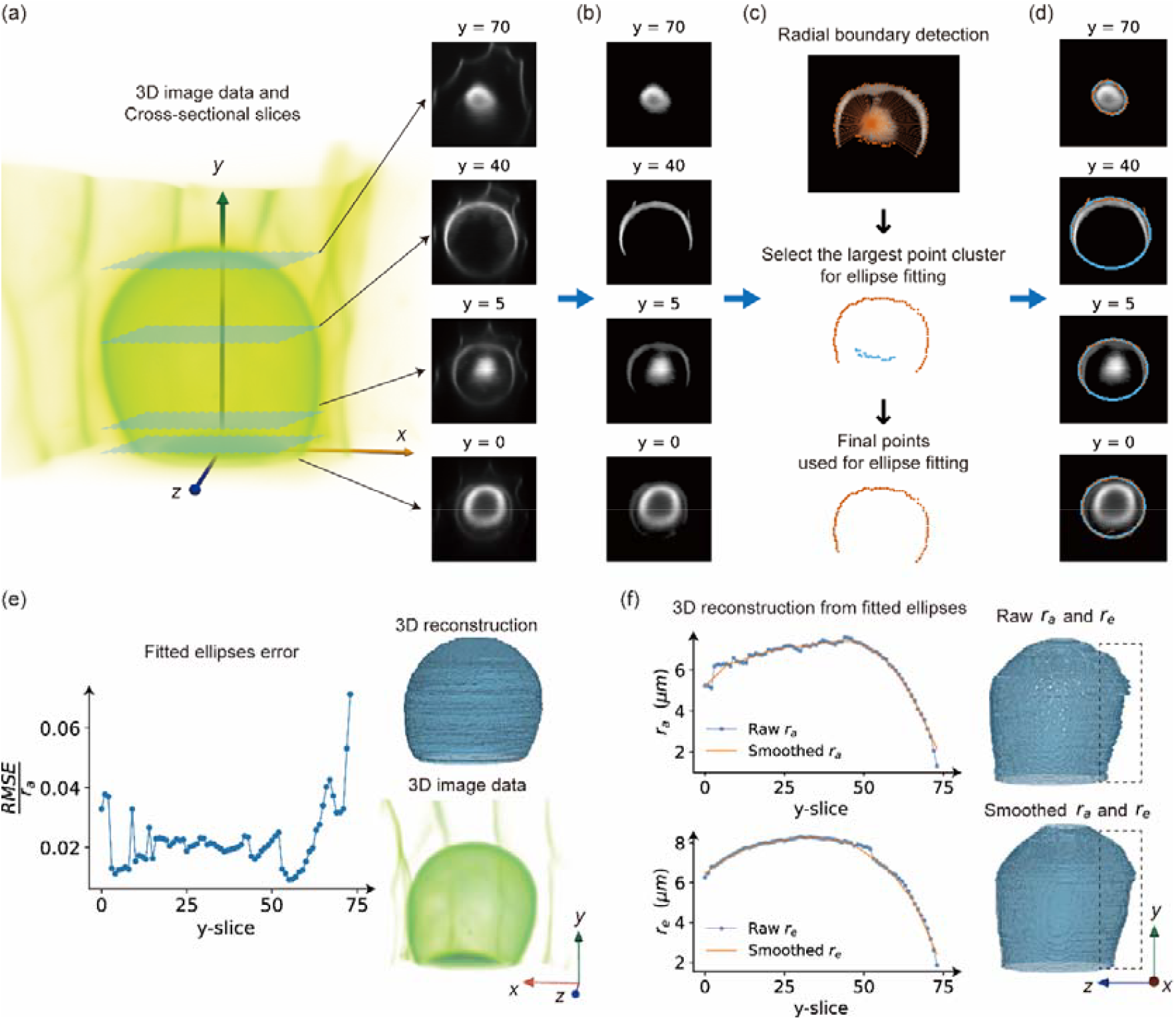
Elliptical cross-section-based reconstruction of 3D cell morphology from normalized voxel data. (a) Extraction of cell cross-sectional contours on planes parallel to the ***xz***-plane from the ***y***-axis-aligned cell after 3D coordinate normalization. (b) Higher intensity region with a threshold to remove background. (c) Radial scanning from the intensity centroid to detect contour points; along each scan line, the outermost pixel exceeding a threshold is selected. When multiple point clusters are detected, the largest cluster is retained as the representative set. (d) Elliptical approximation of the representative point set for each cross-section. (e) Evaluation of ellipse fitting accuracy using the root mean square error (***RMSE***) relative to major axis (***r***_***a***_), confirming consistency with the original intensity-based contour. (f) Lowess smoothing applied to the major and minor axes (***r***_***a***_, ***r***_***b***_) along the ***z***-axis to reduce slice-to-slice variability and obtain a consistent 3D morphological representation.

**Fig. 2.**
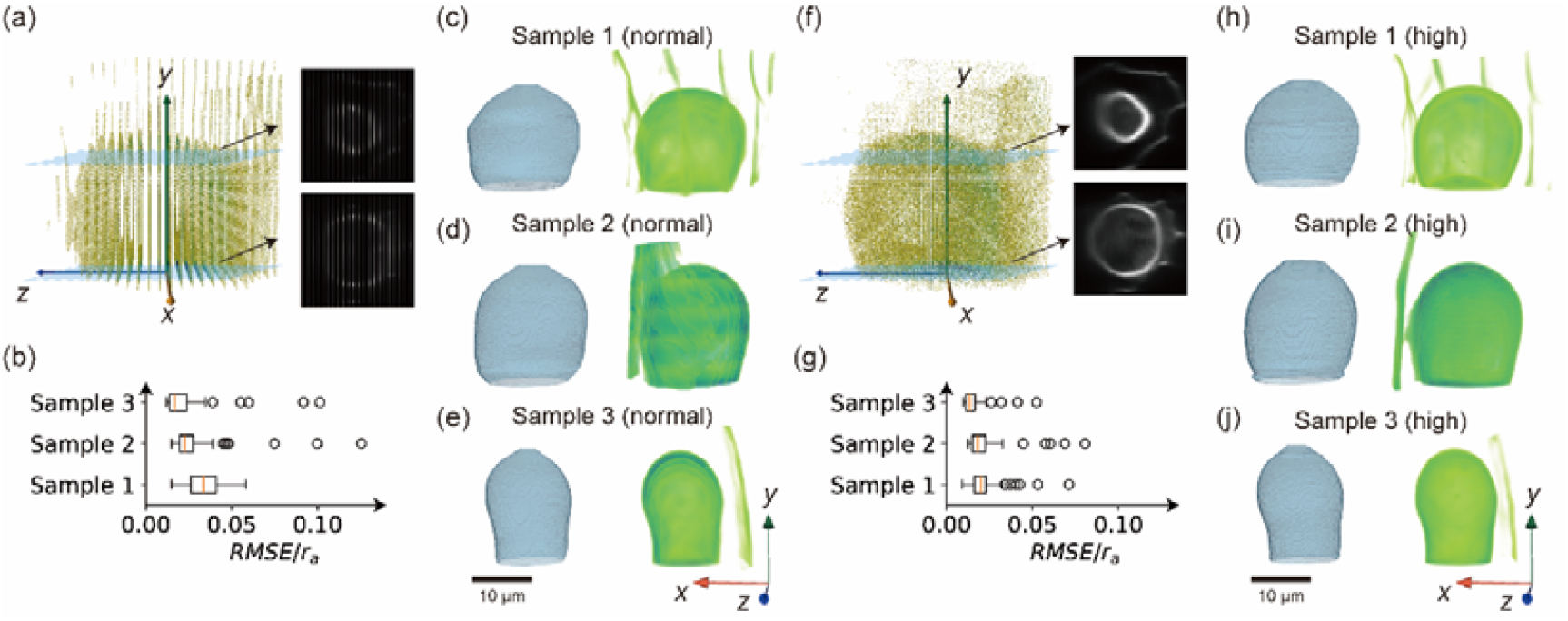
Comparison of the results between the high-resolution images and the normal-resolution images. (a) Normal-resolution image and some cross-sectional images. (b) Relative error of ***RMSE*** / ***r***_***a***_ for the fitting results of the normal-resolution images. (c-e) Obtained 3D reconstruction models for three samples with normal-resolution images. (f) High-resolution image and some cross-sectional images. (g) Relative error of ***RMSE*** / ***r***_***a***_ for the fitting results of the high-resolution images. (h-i) Obtained 3D reconstruction models for three samples with high-resolution images.

### 3. Comparison of the 3D reconstruction between high-resolution images and normal-resolution images

To facilitate 3D reconstruction, the analyses described above used high-resolution images acquired at a 0.2 μm interval along the *Z*-axis (optical axis) (see Methods). Because these imaging conditions were too harsh for long-term observation, we next tested whether our method could also be applied to normal-resolution images acquired at the 1-μm *Z*-interval typically used for embryo live imaging (Fig. 3). Although the difference was evident in the slice density even after coordinate normalization (Fig. 3a and 3f), we confirmed that the resulting ellipse was in a good quality within a relative error of 5% in both cases (Fig. 3b and 3g). In the normal-resolution images, our method successfully corrected sample drift and positional fluctuations during growth, while also compensating for signal loss in dimly illuminated regions, enabling consistent 3D reconstruction (Fig. 3c-e). In the high-resolution images, the same corrections were achieved, and the relative error was further reduced, resulting in a more accurate representation of the original morphology (Fig. 3h-j). These results indicate that high-resolution imaging provides clearer structural information and improves reconstruction accuracy; however, despite a slightly higher relative error caused by sparse images, the normal-resolution images still allow sufficiently reliable 3D reconstruction using the proposed method.

**Fig. 3.**
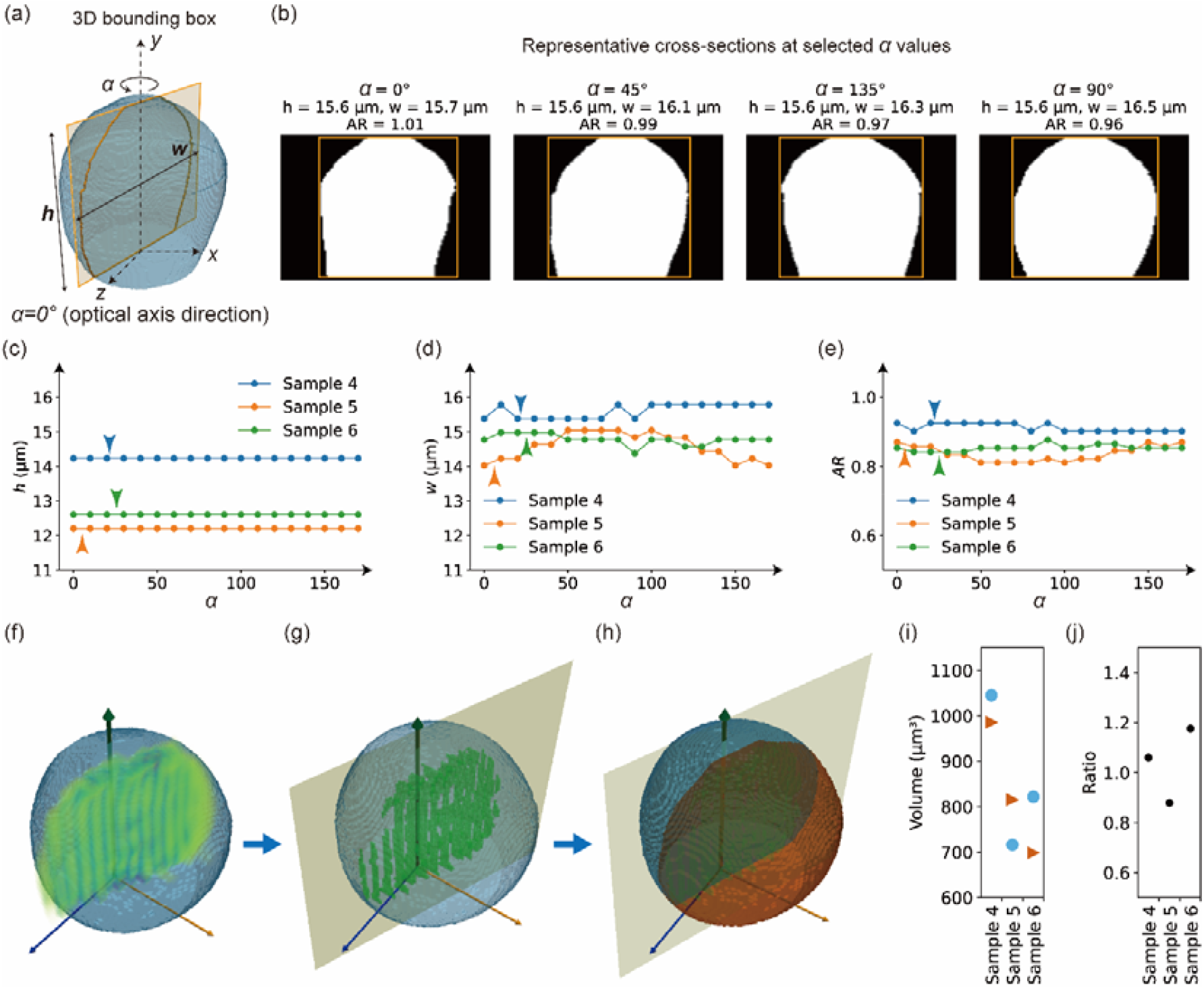
3D bounding-box analysis of the apical cell and volumetric analysis of the two daughter cells. (a) Definition of the bounding box and corresponding rotation angle ***α***. (b) Four representative cell shapes and corresponding bounding-boxes in different directions. (c) Height h of the apical cell in all the directions. Arrow heads show the angle of the actual division plane in data. (d) Width w of the apical cell in all the directions. (e) Aspect ratio (***h / w***) of the apical cell in all the directions. (f) Detection of the point cloud of the newly dividing plane (vertical plane) colored by green points. (g) Fitting a plane for the extracted point cloud. (h) Two daughter cells can be characterized by cutting the apical cell by the fitted plane. (i) Volumes of the daughter cells for three samples. (j) Volumetric ratio of the two daughter cells.

### 4. 3D morphological analysis reveals 3D cell shape in all circumferential directions and morphological traits of divided cells

Finally, the developed Apical3DTip method was used to quantitatively analyze and evaluate the 3D morphology of apical cells just after their vertical cell division (Fig. 4). The reconstructed cell shape is resliced in a specific direction by defining angular orientations *α* on the *xz*-plane and extracting directional cross-sections (Fig. 4a). For each angle, a bounding box enclosing the corresponding projected shape is computed, allowing measurement of its geometric properties in 3D (Fig. 4b). The height *h*, the width *w*, and the aspect ratio of each bounding box (*AR*) were calculated to evaluate morphological characteristics (Fig. 4c-e). Of course, *h* was invariant with viewing direction; neither *w* nor the resulting *AR* (*w*/*h*) showed significant variation with direction, indicating no significant morphological anisotropy at the divided time frame. Using this method, the cell division plane (vertical plane) of the apical cell is also visualized as a point cloud (green) with the coordinate normalization (Fig. 4f). By extracting this point cloud, the orientation of the vertical plane can be analyzed while preserving the original orientation of the apical cell in the standardized coordinate system. The extracted point cloud corresponding to the division plane is then approximated by a fitted plane (Fig. 4g). Subsequently, the apical cell is virtually partitioned by this fitted plane, allowing the volumes of the two daughter cells to be quantified (Fig. 4h-i). Based on these measurements, the volumetric ratio between the two daughter cells can be calculated (Fig. 4j), showing that these samples underwent nearly equal division with the ratio ranging in 0.8-1.2.

These results demonstrate that our method not only enables coordinate normalization and tracking of the morphological changes in apical cells but also allows quantitative analysis of post-divided cellular morphological traits within a common coordinate system, highlighting its broad applicability for 3D morphometric analysis.

## Discussion

We developed Apical3DTip, as a robust pipeline for quantitative 3D analysis of apical cell morphology. By normalizing coordinates based on the cell bottom and optical axis, we established a stable reference frame for consistent comparisons. Elliptic cross-sectional reconstruction analysis provides a compact and noise-reduced representation of 3D geometry. This framework enables quantification of morphological traits around the cell *y*-axis (i.e., the embryo apical-basal axis), and the bottom plane becomes parallel to the *xz*-plane, revealing variations along the embryo radial axis. Extension to 4D analysis may capture spatiotemporal dynamics of shape changes as well as recently developed cellular tip dynamics [9]. The method avoids complex surface reconstruction, making it efficient and suitable for large time-lapse datasets, and provides a foundation for linking cell geometry with developmental processes.

Classical hypotheses of cell division with the minimum surface area rule [4, 5], propose that division orientation is governed by geometric and physical constraints. Although supported by theoretical and experimental studies, quantitative validation in living systems has been limited by the difficulty of capturing dynamic 3D cell geometry. The framework developed here proved useful not only for quantifying cellular isotropy/anisotropy through aspect ratio measurements, but also for detecting the symmetry/asymmetry of cell division based on daughter-cell volume ratios. Future work will evaluate whether cell shape changes reflect mechanical stresses or energetic constraints associated with division plane selection. Our approach therefore provides a platform for directly testing and refining classical models of plant cell division. Furthermore, when combined with KymoTip [10], which enables time-resolved tracking of cell tips, the method in this study may allow accurate 3D tracking of both tip morphology and tip position over time. Together, these features provide a practical framework for routine 3D and 4D analysis of cell morphology and its dynamics. Further combination with recently developed fluorescent probe-based methods for embryo live imaging may also enable quantitative 4D analysis of early embryo patterning in non-model plants, for which methods to generate transgenic lines have not yet been established [14].

Although Apical3DTip was developed for 3D reconstruction of the Arabidopsis apical cell, its applicability is not limited to this specific cell type as long as the cross section is elliptical. The framework is based on geometric features common to tip-growing structures, making it potentially useful for a wide range of biological systems. In plants, pollen tubes and root hairs possess a characteristic cylindrical body with a dome-shaped growing apex [15, 16]. Because Apical3DTip reconstructs and quantifies such geometries in a normalized coordinate system, it can be readily extended to these tip-growing cells. The method may also be applicable to other plant species. For example, the rhizoids of *Marchantia polymorpha* and branching protonemal filaments of *Physcomitrium patens* exhibit a cylindrical organization similar to that assumed by our framework [17, 18]. By enabling quantitative 3D analysis of these structures, Apical3DTip could facilitate comparative studies of polarized growth and morphogenesis, providing a versatile tool for biological shape analysis across diverse species.

Together, these new approaches based on Apical3DTip are expected to reveal fundamental principles underlying plant ontogeny and developmental processes.

## Methods

### A. Plant growth conditions and transgenic lines

All Arabidopsis lines were in the Columbia-0 (Col-0) background. The PM marker (EC1pro:Clover-SYP121, coded as MU2354) consists of the EC1 promoter fragment, the sequence encoding the green fluorescent protein Clover, SYP121, and the NOS terminator in pMDC99 [6]. This EC1pro:Clover-SYP121 was combined with the nuclear marker (EC1pro:H2B-tdTomato and DD22pro:H2B-mCherry, coded as MU1968) [11]. Plants were grown at 20–24°C under continuous light or long-day conditions (16□h light/8□h dark).

### B. Time-lapse observations

The 2PEM live-cell imaging was performed using x40 CFI Apo LWD WI; NA 1.15; Nikon, Japan and a laser-scanning inverted microscope (A1MP or AXMP; Nikon, Japan) equipped with a Ti:sapphire femtosecond pulse laser (Mai Tai DeepSee; Spectra-Physics, Barber Lane Milpitas, CA, USA) or ultrafast tunable laser (InSight X3; Spectra-Physics), respectively as previously described [12-14]. Time-lapse images were taken every 20□min using a 3× zoom as 31 *z*-stacks at 1□μm intervals using A1MP. *Z* stack images were taken with 4.2× zoom as 121 *z*-stacks at 0.2 μm intervals for high-resolution images, or 25 *z*-stacks at 1 μm intervals for normal-resolution images, using AXMP. The obtained images were processed with “Denoise” function in NIS-Elements AR ver. 5.30.02 or 5.42.06 software in A1MP or AXMP, respectively.

### C. Apical3DTip: 3D Coordinate normalization and elliptic fitting of cross sections

#### C-1. Extraction of cell bottom plane

As the cell bottom plane is brighter than the other region, we used the threshold (99.9%) of the fluorescent intensity to remove the region outside of the cell. After this removal, there are still small noise components which are not the bottom plane. To get rid of such small disconnected noise components, we selected the bottom plane which has the largest number of points than the small components. Let the coordinate of the vertices be

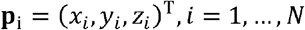

Then the centroid of the bottom plane was calculated as 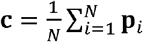, and the centered coordinates were defined as **p**_*i*_ **p**_*i*_ − **c**. We applied Principal Component Analysis (PCA) to the centered point cloud and calculated the eigenvector **n** corresponding to the smallest eigenvalue, which is used as the normal vector of the bottom plane. We normalized the vector as **n**′ = **n /** ∥ **n** ∥. To avoid directional ambiguity, the normal vector was flipped when necessary so that it pointed toward the positive *y*-axis:

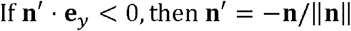

#### C-2. Two-step rotations of the coordinate system

To standardize the orientation of the coordinate system as a positive-*y* axis, we used two types of rotations. This was performed using two-step rotations: first rotation around the *Z*-axis and the second rotation around the *X*-axis. We defined the vector component as **n**′ = (*n*_*X*_, *n*_*Y*_, *n*_*Z*_)^T^. The first rotation angle was defined as

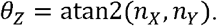

The corresponding Z-axis rotation matrix was

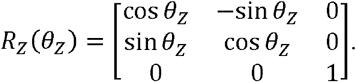

After *Z*-axis rotation, the coordinate system is rotated into a new coordinate system (*X*_1_, *Y*_1_, *Z*_1_) with the normal vector as 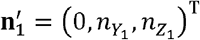. In this new coordinate system, the *X*_1_-component of the normal vector becomes zero, and the vector lies in the *Y*_1_ *Z*_1_-plane. The second rotation was performed around the *X*_1_-axis. The rotation angle was defined as

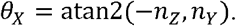

Then the *X*_1_-axis rotation matrix was

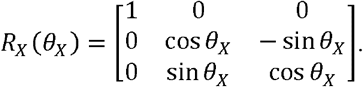

The total rotation matrix was *R* =*R*_X_ (*θ*_*X*_) *R*_*Z*_ (*θ*_*Z*_). After rotation, the normal vector satisfies

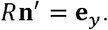

**Fig. S1.**
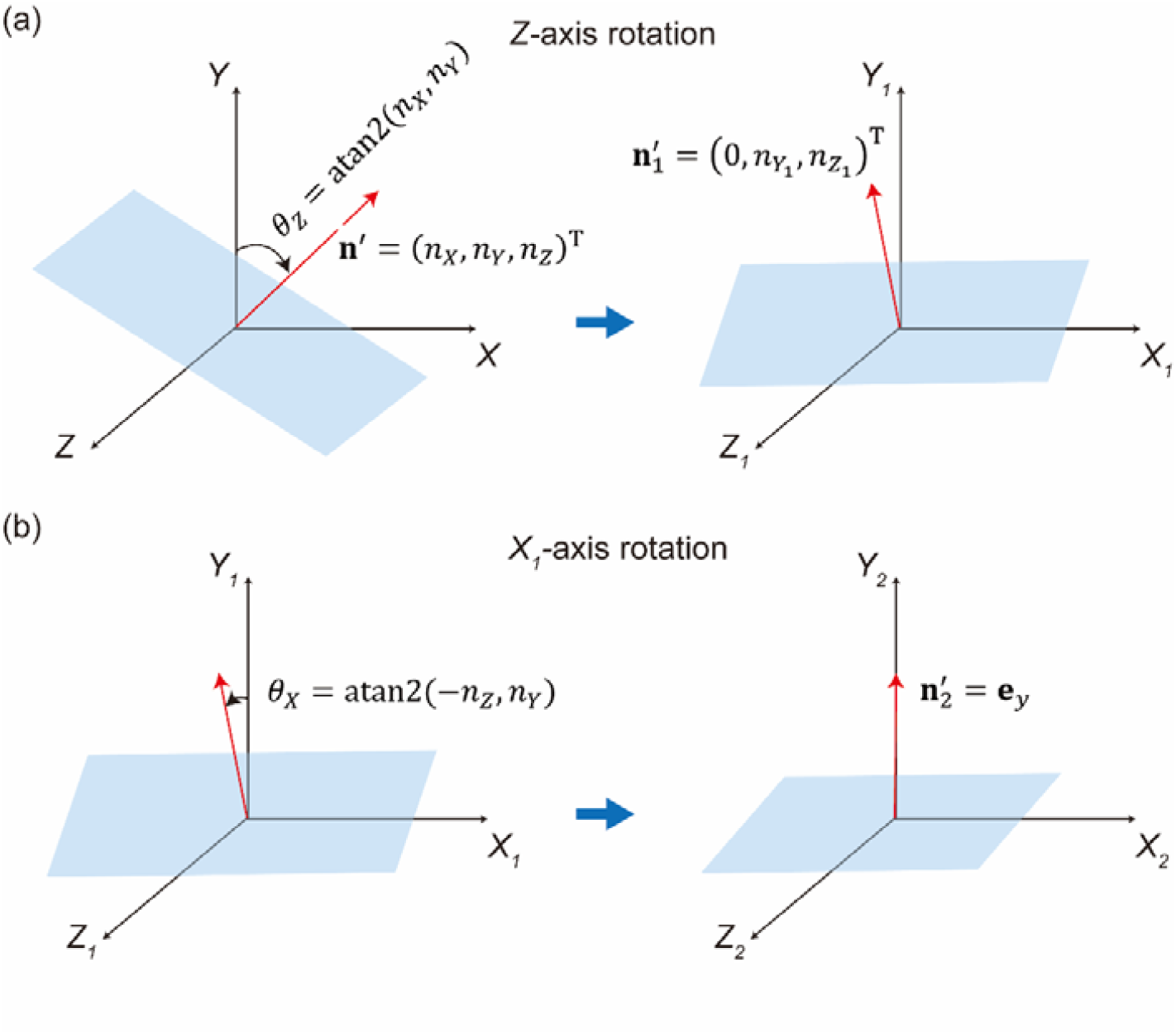
Schematic illustrations of two types of rotations of the original coordinate system. (a) *Z*-axis rotation is performed by the rotation angle *θ*_*Z*_ around the *Z*-axis and the resulting coordinate system becomes (*X*_1_, *Y*_1_, *Z*_1_). (b) *X*_1_-axis rotation is performed by the rotation angle *θ*_*X*_ around the *X*_1_-axis and the resulting coordinate system becomes (_2_ **= *x***, *Y*_2_ = *y, Z*_2_ = *z*) where the *y*-axis coincides with the vertical axis perpendicular to the bottom plane.

#### C-3. Radial search and DBSCAN clustering

With the above procedures, the cell coordinate system is properly standardized with the bottom plane as the *xz*-plane. We resliced the cell along the *y*-axis to obtain the contour on the *xz*-plane as a function of *y*. There were a few candidate contour points so that we used radial searching on each resampled the points on the *xz*-plane. A slice-dependent threshold was set to the 60 percentiles of non-zero intensities. The center of searching was defined as the centroid of the extracted points above the slice-dependent threshold. From this center, we performed radial search with 120 divisions, and the fluorescent points along each sector were detected. Since the candidate boundary points often included outliers, DBSCAN clustering (python library) was applied to identify the dominant spatially connected point group. We identified the DBSCAN groups points based on local point density and labeled points with low-density regions as outliers. The neighborhood radius was set to *ϵ* = 5 *μm*, and the minimum number of neighboring points required to form a cluster was set to 5. The largest cluster was regarded as the final contour point set.

#### C-4. Ellipse fitting and calculation

An ellipse was then fitted to the final contour point set using the OpenCV ellipse-fitting algorithm. The fitted ellipse was expressed as

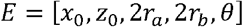

where (*x*_0_, *z*_0_) is the center, 2*r*_*a*_ and 2*r*_*b*_ are the major- and minor-axis lengths, and *θ* is the rotation angle (Fig. S2a). Each contour point **q**_*i*_ **=** (*x*_*i*_, *z*_*i*_)^T^, was first translated relative to the fitted ellipse center and then rotated by − *θ* (Fig. S2b),

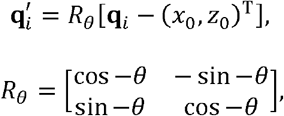

so that the ellipse axes became aligned with the axes of the coordinate system (Fig. S2c). This transformation enabled radial distances between the observed contour and the fitted ellipse to be calculated in a common ellipse-aligned coordinate system with polar coordinate (*r*_pred,*i*_ Φ_*i*_).

The point angle and ellipse radius were as follows.

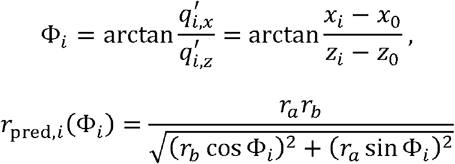

The observed radial distance in data was

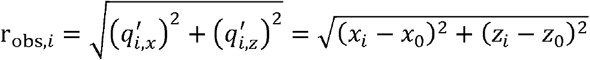

The fitting error was calculated as

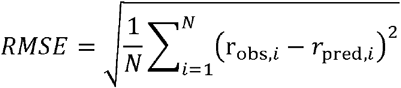

**Fig. S2.**
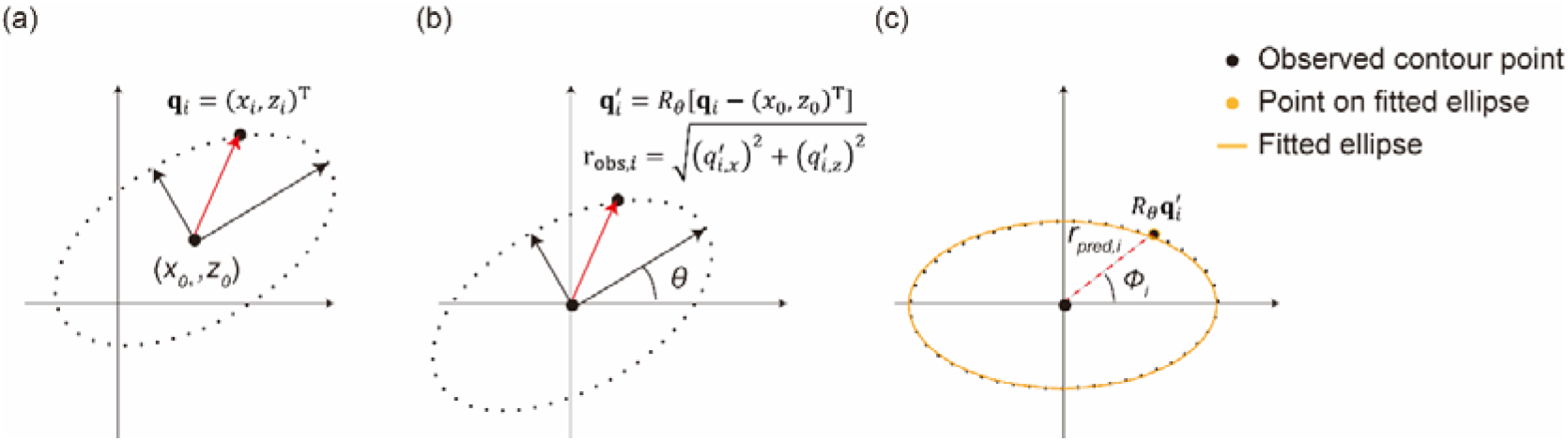
Schematic illustrations of elliptic fitting processes. (a) Data points *q*_*i*_ = (*x*_*i*_, *z*_*i*_) are fitted by the ellipse with the center (*x*_0_, *z*_0_). (b) Translation is performed to locate the ellipse center at the center of the coordinate system, and rotation is performed to align the major and minor axes of the ellipse coincide with the axes of the coordinate system. (c) Calculation of the fitting error. The *RMSE* is calculated between the aligned observed contour point and the point on the fitted ellipse.

#### C-5. LOWESS smoothing

To reduce slice-to-slice fluctuations along *y*-axis, the major and minor axis lengths (2*r*_*a*_, 2*r*_*b*_) of the fitted ellipse were smoothed along the *y*-axis using locally weighted scatterplot smoothing (LOWESS). The LOWESS smoothing was implemented in Python using the *lowess* function from *statsmodels*.*nonparametric*.*smoothers_lowess* module. The smoothing fraction parameter was set to *η* = 0.2, meaning that 20% of the neighboring points were used for each local regression.

#### C-6. Volumetric analysis of two daughter cells

The vertical plane is defined as the newly formed division plane of the apical cell after the first cell division at the bottom. The vertical plane was assigned to the set of points corresponding to the common boundary of the daughter cells. Let these points be

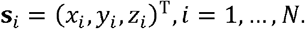

A plane in 3D space can be described by a unit normal vector **n**_0_ and an offset *e*:

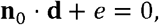

where **d** denotes an arbitrary point on the fitted plane. The best-fit plane was obtained by minimizing the squared orthogonal distances from the interface points to the fitted plane:

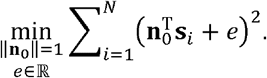

In practice, the fitted vertical plane was estimated by PCA. First, the centroid of the interface points was calculated:

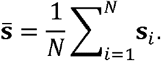

The eigenvector corresponding to the smallest eigenvalue of the interface points was used as the normal vector of the fitted vertical plane:

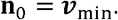

Because the fitted plane passes through the centroid s, the offset e was calculated as

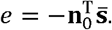

Then, the fitted plane was represented as

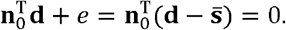

## Availability of data and materials

The code for Apical3DTip, along with all associated datasets, is available on Github: https://github.com/blues0910/Apical3DTip.

Apical3DTip is also available as an ImageJ plugin: https://github.com/YusukeKimata-Moo/Apical3DTip.

## Ethics approval and consent to participate

Not applicable.

## Consent for publication

Not applicable.

## Authors contributions

Z.K., T.N., S.T. and M.U. designed the study. Y.H., Y.I. and H.M. performed the experiments. Z.K. and T.N. performed 3D reconstruction. Z.K., T.N. and Y.K. contributed materials/analysis tools. Z.K., T.N., S.T. and M.U. wrote the manuscript. All authors reviewed the manuscript.

## Acknowledgements

The authors thank Koichi Fujimoto and Naoya Kamamoto (Hiroshima University), and Takumi Higaki (Kumamoto University) for helpful discussions.

